# Cyclic-di-GMP controls Type III effector export and symptom development in *Pseudomonas syringae* infections via the export ATPase HrcN

**DOI:** 10.1101/2025.07.16.665098

**Authors:** D. Ward, R.H. Little, C.M.A. Thompson, J.G. Malone

## Abstract

*Pseudomonas syringae* is a destructive bacterial pathogen that infects a wide variety of plants. Following apoplastic entry, *P. syringae* uses its type 3 injectisome (T3I) to secrete host-specific effectors into the cytoplasm, enabling tissue wetting and immune suppression and leading to bacterial proliferation, chlorosis and necrosis. *P. syringae* strains encode dozens of highly specialised effectors, whose composition defines strain specificity and host range. Effective plant infection depends on the tight temporal and hierarchical control of effector delivery through the T3I. Effector secretion is driven by HrcN, an ATPase complex that interacts with the base of the T3I and is essential for plant infection. HrcN binds specifically to the bacterial signalling molecule cyclic-di-GMP, although the impact of binding on T3I function and *P. syringae* virulence is currently unknown. To address this, we examined the influence of mutating the predicted cyclic-di-GMP-*hrcN* binding site on plant infection and effector secretion. Despite maintaining effective bacterial proliferation in *Arabidopsis thaliana* leaves, two *hrcN* mutants showed severely compromised disease symptoms, a phenotype linked to reduced translocation of a specific subset of T3I-effectors, with HopAA1-2 particularly important for symptom development. We propose that cyclic-di-GMP binding may represent a novel regulatory mechanism for effector secretion during bacterial infections.

## Introduction

*Pseudomonas syringae* is a diverse species complex of Gram-negative foliar plant pathogens, comprising over 100 distinct pathovars (1) with the ability to infect a wide variety of plants, including almost all economically important crops (2). *P. syringae* is a widely used model organism for the molecular analysis of plant–microbe interactions (2, 3), with the tomato pathogen *P. syringae* pv. *tomato* DC3000 (*Pto*) particularly well-studied due to its ability to infect the model plants *Arabidopsis thaliana* and *Nicotiana benthamiana* (4, 5). Upon entry into the apoplast through surface wounds and leaf stomata, *P. syringae* infections proceed via tissue wetting accompanied by rapid bacterial proliferation, then chlorosis and eventual tissue necrosis (6).

A key organelle in the infection process is the type 3 injectisome (T3I), with a functional T3I necessary for plant infection and apoplastic proliferation (7). This large, multiprotein complex spans the inner and outer membranes, connecting the bacterial cytoplasm to that of its host and enabling the translocation of host-specific protein effectors into the plant cytoplasm (8). The core T3I components share a high level of structural conservation and a close evolutionary lineage with the secretory system of the bacterial flagellum (9). T3I assembly is tightly regulated, with extracellular subunits exported in sequence through the central pore of the lengthening organelle (10).

Recruitment and secretion of type 3 effectors (T3Es) proceeds via the recognition of highly variable secretion signals at the effector N-terminus (11, 12) in a chaperone-dependent or independent manner (13). Effector proteins are then recruited to a sorting platform at the base of the T3I (13, 14). This sorting platform consists of a C-ring (HrcQ and HrpD), export apparatus (HrcRST, HrcU and HrcV) and an ATPase complex (HrpE and HrcN) (15, 16). The HrcN ATPase is a central component of T3I function, whose deletion almost entirely abolishes bacterial cytotoxicity (17).

HrcN forms a dodecamer at the base of the sorting platform (18) and controls substrate entry into the export gate by ensuring that ATP hydrolysis occurs in concert with secretion. It also ensures efficient substrate unfolding in the export channel (19–21). T3I ATPase monomers comprise a central ATPase domain flanked by an N-terminal domain containing a 5 stranded β-sheet and a predominantly α-helical C-terminal domain (22–24). The cryo-EM structure of the closely related *Escherichia coli* ATPase EscN reveals an asymmetric multimer, suggesting a rotational catalysis mechanism similar to that of F1 ATPases (22).

T3Es encoded by *P. syringae* pathovars fulfil diverse roles in plant infection, facilitating bacterial colonisation while repressing the host immune response (1, 25). T3Es can trigger apoplastic wetting (26), manipulate host NAD(+) metabolism (27), suppress plant immune responses (28–32) or interfere with other traits including organelle function, cell development, nutrient distribution and metabolism (33). *P. syringae* strains typically contain 15-35 well-expressed T3Es (∼36 in *Pto* (34)), drawing from at least 58 distinct effector families (35, 36). The effector repertoire of an individual strain is highly specialised towards colonisation of its host plant, with these differences one of the main factors defining strains and pathovars (37).

Effective plant infection depends on the tight temporal and hierarchical control of T3E delivery (38). For example, effectors enabling apoplastic wetting are deployed in the initial stages of infection (26), while early- and late-stage immune responses are targeted by different T3Es. AvrPtoB targets the PAMP immune receptor FLS2 for degradation (39), while HopM1 disrupts late-stage immunity by inducing degradation of the vesicle trafficking-related protein AtMIN7 (40). Leaf chlorosis and necrosis are apparently enabled by targeted translocation of a few key T3I effector proteins, rather than with a larger fraction of the effector protein repertoire (41–43). Coordinated effector delivery to the T3I is enabled by specific *hrp*-associated (*hpa*) chaperones (44). Furthermore, T3I deployment and activity are also subject to extensive regulatory control (10, 45).

Our previous research showed that the ubiquitous dinucleotide signalling molecule cyclic-di-GMP (cdG) binds specifically to the ATPase proteins of several bacterial secretory systems, including the flagellar protein FliI and HrcN from *Pto* (46). CdG is associated with virulence and infectious persistence in many human and plant pathogens (47) and exerts its effects by binding to diverse protein and RNA targets, which in turn control an array of bacterial traits (48). In *P. syringae*, cdG tightly controls the transition between motile, unicellular and communal, biofilm-forming lifestyles (49), with T3I-mediated virulence among the regulated traits (45). CdG binding to HrcN-like ATPases appears to be widespread, with examples seen with type II, III, IV and VI secretory systems from various species (46, 50–52). At this stage, however, the impact of cdG-HrcN binding on T3I activity and *Pto* virulence is unknown.

To examine this relationship in more detail, we produced mutants in several *Pto hrcN* residues predicted to play a role in cdG-binding, and tested their impact on plant infection, protein function and T3I effector secretion. Two *hrcN* mutants, *G176A* and *G311A*, showed severely compromised visual disease symptoms, despite maintaining effective bacterial colonisation of *A. thaliana* Col0 leaves. This asymptomatic phenotype could be reversed by disrupting *Pto* intracellular cdG levels, and depends on a functional plant immune response. Using *cyaA*-fusion effector translocation assays (53), we observed significantly reduced translocation of a subset of T3I-effectors in the *G176A* mutant compared to WT *Pto*, with the asymptomatic phenotype linked to impaired HopAA1-2 secretion in particular.

## Results

### Specific HrcN mutations ameliorate disease symptoms upon *P. syringae* leaf infection

In our previous work we showed that cdG binds specifically to the HrcN ATPase, with dinucleotide binding sites predicted to lie at the interface between two monomers within the larger dodecameric complex at the base of the *P. syringae* T3I (54). At this stage, however, the biological function of this interaction is unknown. To address this, we first highlighted several residues predicted in our original study to contribute to the HrcN cdG binding site (54) on an AlphaFold 3 model of *P. syringae pv. tomato* DC3000 (*Pto*) HrcN (Fig 1A). The predicted positions of these residues was broadly in agreement with our earlier work (54), with the predicted cdG binding sites lying at the interface between HrcN subunits, with each residue either contributing, or coordinating residues that contribute, to the binding pocket (Fig 1B).

**Fig 1A:**
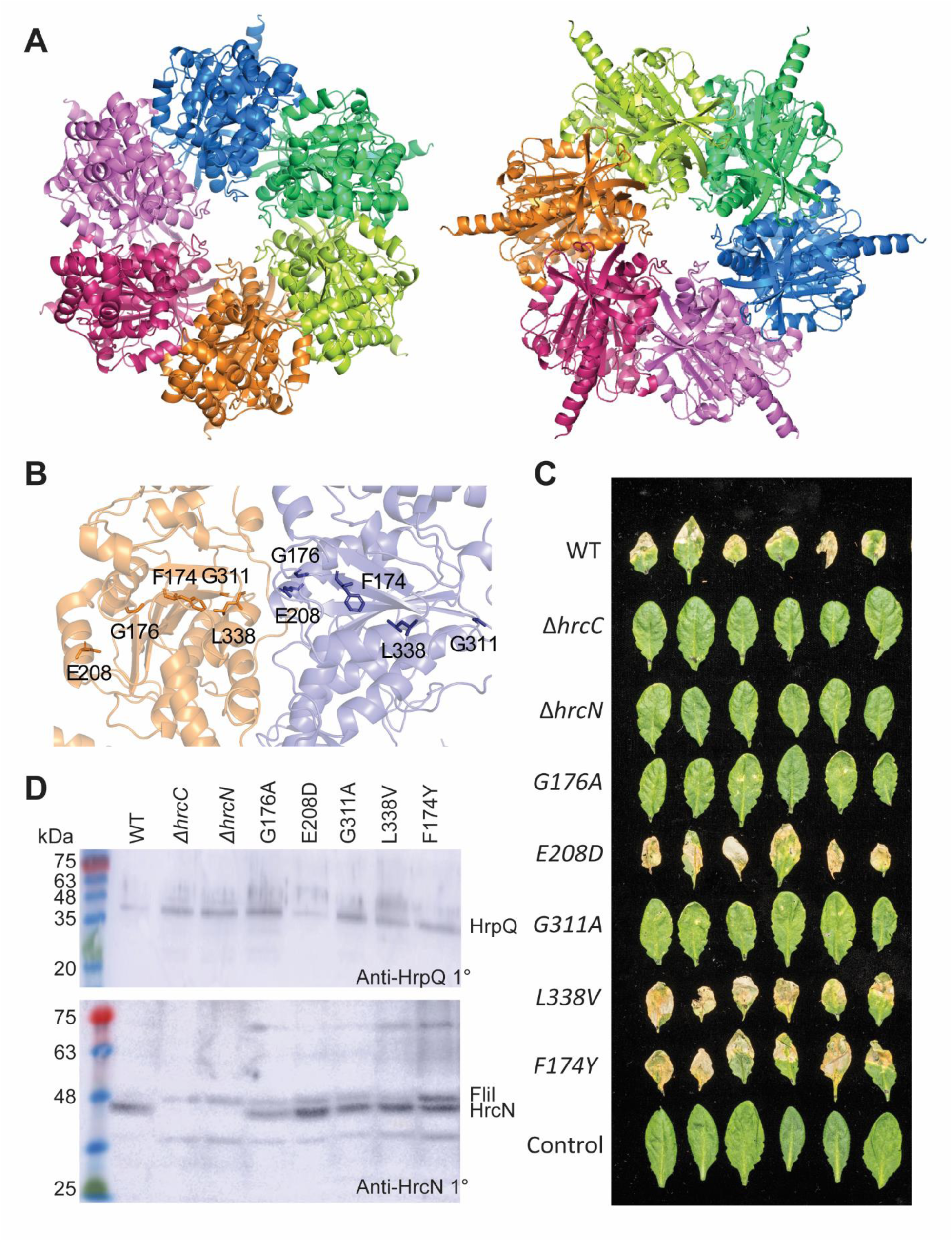
AlphaFold3 structural predictions for hexameric HrcN complexes. Left: C-terminus facing; Right: N-terminus facing. **1B:** Predicted cdG binding site residues mutated in this study, highlighted on the HrcN structure. Note that each binding site is predicted to incorporate residues from two adjacent subunits. **1C:** *A. thaliana* Col-0 disease phenotypes at 6 days post-infiltration infection with *Pto* DC3000 strains containing deletions in *hrcC/hrcN*, alongside mutations leading to residue substitutions in the predicted *hrcN* cdG binding site and an uninfected control. Six representative leaves from four plants are shown in each case. **1D:** Anti-HrpQ (top) and Anti-HrcN (bottom) Western blots of *A. thaliana* Col-0 leaf tissue infected with *Pto* DC3000 containing *hrcC/hrcN* mutant alleles. Infected leaf tissue was collected three days post-infection. The locations of HrpQ (present in all samples) and HrcN (absent from Δ*hrcC* and Δ*hrcN*) are indicated. FliI appears as a contaminating band in the Anti-HrcN blot, with all three protein identities confirmed by mass spectrometry.

Next, we constructed a series of mutants, introducing conservative substitutions of each selected residue into *Pto hrcN*. We then tested the capacity or each mutant to infect Col0 *Arabidopsis* plants, relative to WT *Pto* and Δ*hrcC* & Δ*hrcN* negative control strains. Bacterial load was measured for each strain at 2 and 3 days post-leaf infiltration. The Δ*hrcC*/Δ*hrcN* negative controls showed little increase in bacterial load from the initial inoculum, consistent with the loss of T3I function (S1A Fig). Meanwhile, a 3-4 log increase in load was observed for WT *Pto* and all five tested mutants.

Six days post-bacterial infiltration with WT *Pto*, infected leaves displayed systemic chlorosis and tissue necrosis, in contrast to the negative controls where no symptoms of infection were observed. Strikingly, however, while the disease symptoms for *hrcN^E208D^*, *hrcN^L338V^* and *hrcN^F174Y^* infections did not differ noticeably from WT *Pto*, minimal disease symptoms were observed for the *hrcN^G176A^* and *hrcN^G311A^* mutants, with little visible chlorosis or necrosis in either case (Fig 1C). Observed differences in leaf discolouration were quantified using an average pixel intensity calculation applied to each leaf image using ImageJ (version 1.52a), with highly significant differences in the proportion of discoloured leaf tissue observed between WT *Pto, hrcN^G176A^* and *hrcN^G311A^* (S1B Fig).

Western blotting of infected leaf tissue 3 days post-infiltration confirmed the presence of HrcN in all samples except for Δ*hrcC*, Δ*hrcN* and the uninfected plant tissue control. Likewise, Western blotting detected the inner membrane structural protein HrpQ in all samples except for the uninfected negative control, confirming that *hrcC* and *hrcN* mutations do not impact the deployment of HrpQ (Fig 1D).

We then complemented the *hrcN/hrcC Pto* mutations by integrating a WT copy of *hrcN* (or *hrcC*) under the control of its native promoter into the neutral chromosomal *att*::Tn*7* site of each strain. Full recovery of tissue chlorosis and necrosis was observed upon leaf infection with the complemented *hrcN^G176A^* and *hrcN^G311A^* strains, suggesting that the asymptomatic infection phenotype resulted from the *hrcN* mutation in each case. Δ*hrcC* and Δ*hrcN* also displayed WT infection phenotypes when complemented, with no differences in infection phenotypes observed for the other complemented mutants (S1C-D Fig).

### The G176A mutation does not compromise HrcN ATPase activity or cdG binding

To probe the relationship between cdG-binding and HrcN function in more detail, we next expressed and purified N-terminal His6-tagged HrcN alongside its G176A and G311A variants. All three protein variants were unstable and rapidly precipitated from solution, with the G311A variant performing most poorly. We therefore decided to restrict our biochemical analysis to the G176A variant. Both WT HrcN and HrcN^G176A^ maintained ATPase activity, in line with our previous findings (54) (Fig 2A). Likewise, we confirmed cdG binding to WT HrcN using DRaCALA assays (55) with fluorescent-tagged cdG (Fig 2B). Curiously, the HrcN^G176A^ variant also bound cdG in these experiments (Fig 2B-C), in contrast to earlier results seen for *P. fluorescens* FliI (54). CdG binding to both HrcN variants was specific; fluorescent cdG binding could be disrupted by the addition of unlabelled cdG, but not by a range of other nucleotides (Fig 2C). Furthermore, neither HrcN variant could bind the dinucleotide cyclic-di-AMP (cdA, Fig 2D).

**Fig 2A:**
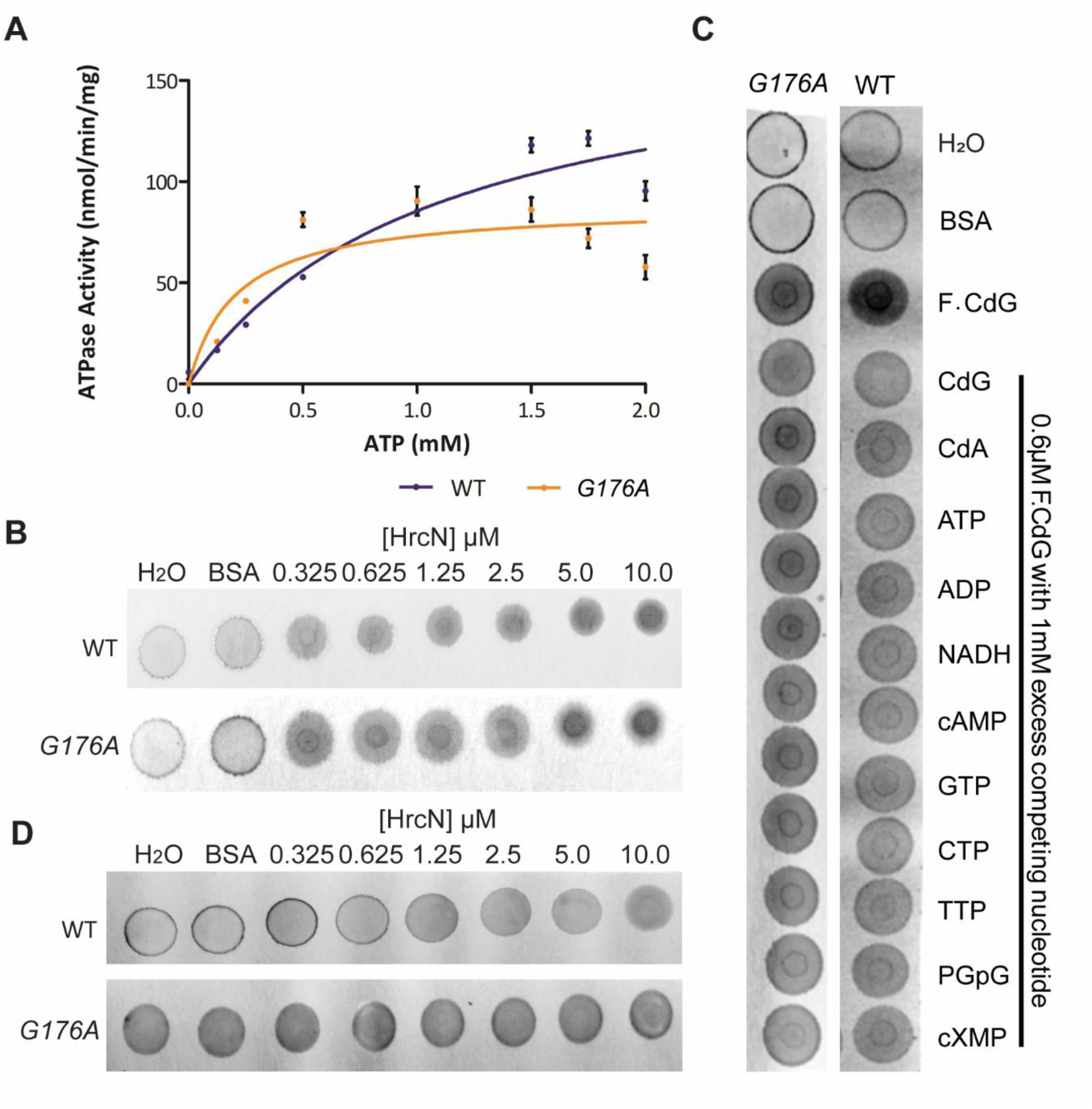
ATPase activity demonstrated for purified WT HrcN and HrcN^G176A^ using a PK/LDH ATPase assay. This reaction has been plotted as a Michaelis-Menten saturation curve where [HrcN] is 1 µM. Error bars show standard error of the mean. **2B:** Fluorescent-CdG DRaCALA assays with increasing concentrations of purified Δ1-18 WT HrcN (top) and Δ1-18 HrcN^G176A^ (bottom). Fluorescent-CdG is included at a fixed 0.6 µM final concentration. **2C:** Competition DRaCALA assays with excess unlabelled nucleotides. Δ1-18 WT HrcN / HrcN^G176A^ and fluorescent-CdG are present at fixed final concentrations of 1.25 µM and 0.6 µM, respectively. Non-labelled competitor nucleotides, as indicated, were added to a final concentration of 1mM. **2D:** Fluorescent-CdA DRaCALA assays with increasing concentrations of purified Δ1-18 WT HrcN and Δ1-18 HrcN^G176A^. Fluorescent-CdA is included at a fixed 0.6 µM final concentration. **B-D:** Control samples contain either H_2_O or 10 µM BSA in place of HrcN, as indicated.

### The *hrcN^G176A^* and *hrcN^G311A^* infection phenotypes are linked to bacterial cdG levels and plant immunity

To investigate the *hrcN^G176A^* and *hrcN^G311A^* asymptomatic infection phenotypes in more detail, we conducted additional rounds of leaf infection with strains over-expressing the well-characterised cdG synthase and phosphodiesterase genes *wspR^19^* and *bifA* (49, 56), producing strains with elevated and suppressed cdG levels, respectively. CdG overproduction by WspR^19^ fully recovered tissue chlorosis and necrosis in both *hrcN^G176A^* and *hrcN^G311A^* infections (Fig 3A, S2A-B Fig), suggesting that excess cdG is able to override the effects of the *hrcN* mutation in each case. Curiously, symptom recovery was also observed for low-cdG, *bifA-*expressing strains, albeit only partially in the case of *hrcN^G176A^* (Fig 3B, S2C-D Fig). Disruption of cdG levels did not produce noticeable effects on infections with any of the other tested mutants or control strains.

**Fig 3A-B:**
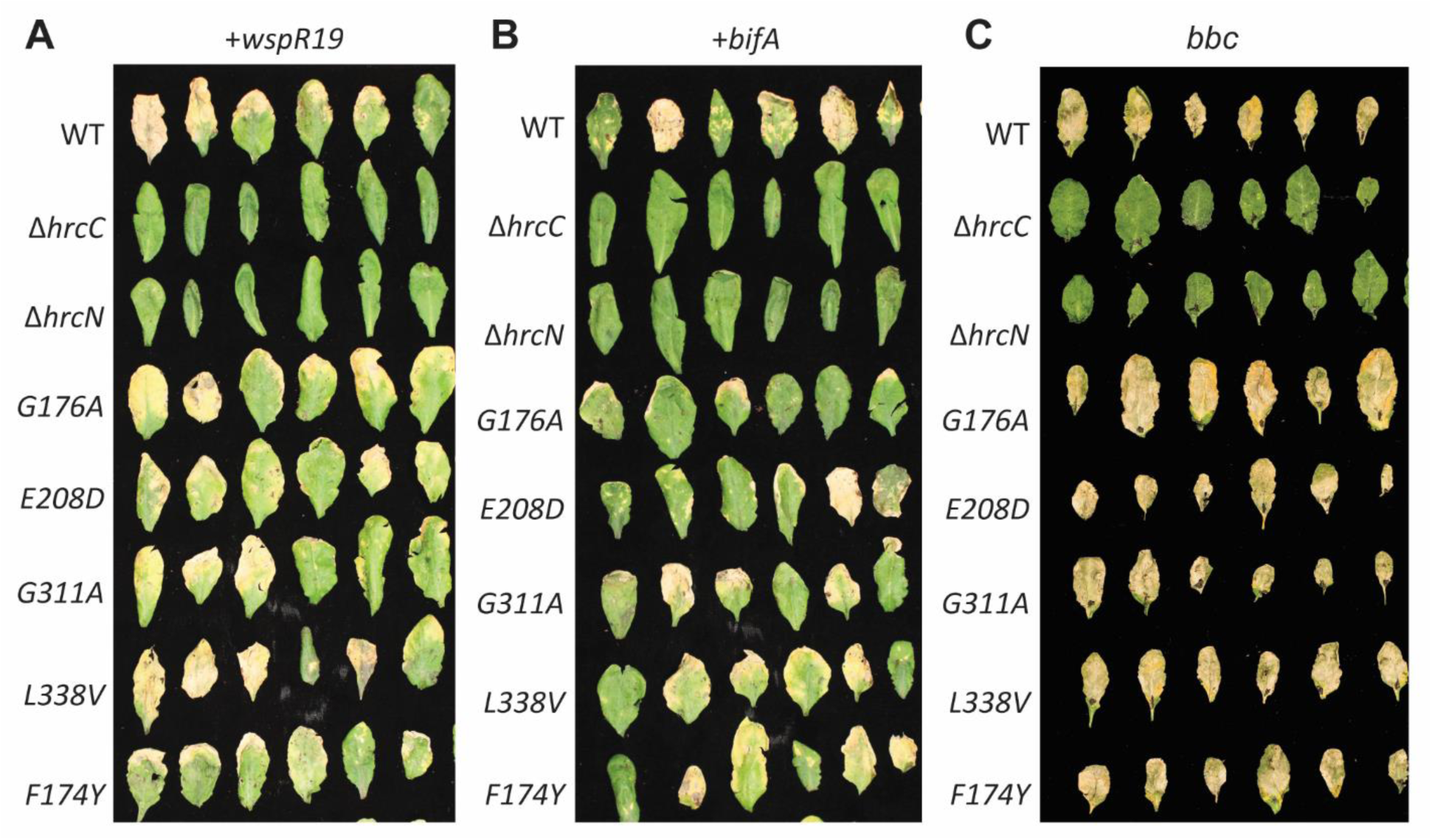
Visual disease phenotypes of *Pto* DC3000 infiltrated *A. thaliana* Col-0 leaves, 6 days post-infection. *Pto* strains harbour mutant *hrcC*/*hrcN* alleles as indicated, alongside **A:** pBBR4-*wspR19* (inducing high cdG levels) and **B:** pBBR4-*bifA* (inducing low cdG levels). **3C:** Disease phenotypes of *Pto* DC3000 infiltrated *A. thaliana bak1-5,bkk-1,cerk1* (*bbc*) leaves, 6 days post-infection. *Pto* strains harbour mutant *hrcC*/*hrcN* alleles as indicated. Leaves were taken at random from three independent plants in each case.

We next examined the impact of *hrcN* mutation on infections of immune compromised plants. Infection of Col0 *bbc* plants; missing the *bak1-5, bkk-1*, and *cerk1* immune receptor genes (26), with WT *Pto* and all five *hrcN* mutants led to strong infection phenotypes, in contrast to the Δ*hrcC*, Δ*hrcN* and water controls (Fig 3C, S3A-B Fig). Similar results were seen with Col0 *fec* plants (with *fls2, efr* and *cerk1* immunity genes deleted (57)). Here, strong infection phenotypes were observed for WT *Pto*, *hrcN^G176A^*, *hrcN^E208D^* and *hrcN^G311A^* in contrast to the Δ*hrcC* negative control (S3C-E Fig). These data suggest that the asymptomatic phenotype of *hrcN^G176A^* & *hrcN^G311A^* mutants may stem from reduced secretion of effector proteins that target the *Arabidopsis* immune system.

### *Pto hrcN^G176A^* and *hrcN^G311A^* mutants display compromised T3I effector secretion

To probe the relationship between the *hrcN^G176A/G311A^* infection phenotypes and effector secretion, we constructed CyaA-fusion vectors for 23 randomly selected *P. syringae* effectors (58). These effector fusions are unfolded until they translocate into plant tissue, where translocation can be detected using ELISA assays for cAMP synthesis (58). The vectors were transformed into WT *Pto,* Δ*hrcN, hrcN^G176A^* and *hrcN^G311A^*, plants were infected, and relative effector translocation measured in each case (Fig 4A, B). Strikingly, both *hrcN^G176A^* and *hrcN^G311A^* showed substantial differences in the extent of translocation relative to WT *Pto* for multiple effectors. For *hrcN^G311A^*, five effectors showed >2-fold elevated translocation, while four showed levels less than half those of WT *Pto*. For *hrcN^G176A^*, no effectors showed >2-fold elevated translocation, while nine showed >2-fold reduced translocation. No effector translocation was observed for any Δ*hrcN* strain, as expected.

**Fig 4A:**
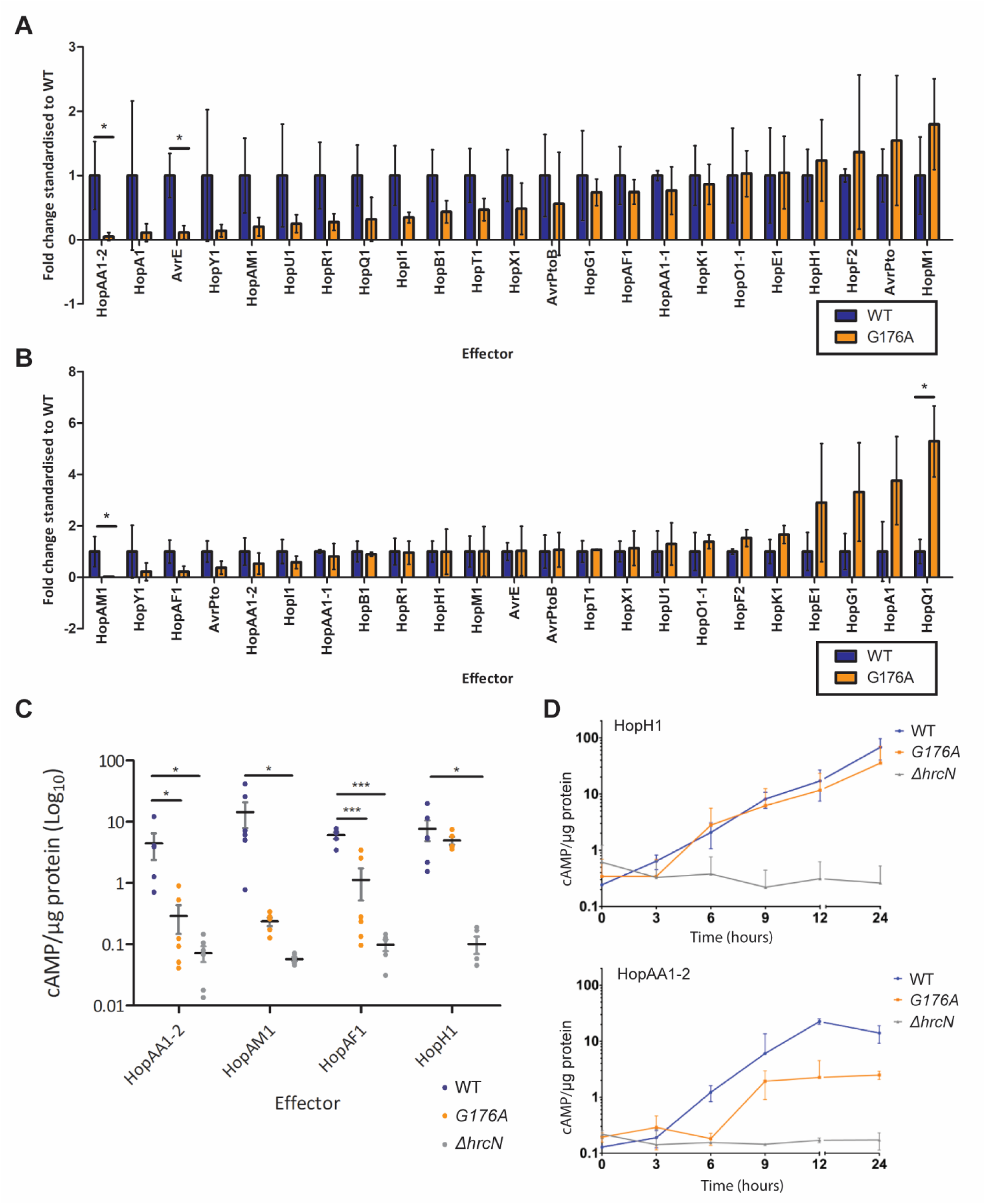
*Pto hrcN^G176A^* effector-CyaA translocation rates shown relative to WT *hrcN*. **4B:** *Pto hrcN^G311A^* effector-CyaA translocation rates shown relative to WT *hrcN*. In each case, values above 1 indicate an increase in translocation for a given effector in the *hrcN^G176A^* /*hrcN^G311A^* mutant compared to WT, values below 1 show a decrease (n = 3). **4C:** Effector-CyaA fusion reporter assays for key *Pto* effectors showing cAMP/µg protein in *Pto* DC3000 infected *A. thaliana* Col-0 leaves 6 hours post-infection, where cAMP/µg protein directly correlates with effector translocation. In each case, asterisks (*) denote statistical significance (2-sample t test, n = 6) compared to the mean WT value, where ‘*’ = p ≤ 0.05 and ‘***’ = p ≤ 0.01. **4D:** Effector-CyaA fusion reporter assays showing cAMP/µg protein in *Pto* DC3000 infected *A. thaliana* Col-0 leaves over 24 hours post-infection. Top graph: HopH1 effector translocation, bottom graph: HopAA1-2. *Pto* strains harbour mutant *hrcN* alleles as indicated.

We next selected four effectors for more detailed analysis. HopAA1-2 and HopAM1 showed the greatest translocation reductions in *hrcN^G176A^* and *hrcN^G311A^* respectively and were both strongly reduced in *hrcN^G176A^*, while HopAF1 showed strongly reduced translocation in *hrcN^G311A^* and mildly reduced translocation in *hrcN^G176A^*. HopH1 showed little difference in translocation across any of the tested strains and was included as a negative control. Effector translocation assays with these four proteins were then conducted in WT *Pto,* Δ*hrcN* and *hrcN^G176A^*. In line with our previous findings, absolute levels of effector secretion from *hrcN^G176A^* were significantly lower than WT *Pto* for HopAA1-2, HopAM1 and HopAF1, while no significant difference was observed for HopH1 (Fig 4C). When measured over the course of 24 hours post-infection, the onset of HopAA1-2 translocation was delayed by approximately 3 hours in the *hrcN^G176A^* mutant compared to WT *Pto*, and occurred to a significantly reduced extent. No differences were seen for HopH1 translocation over the same time period (Fig 4D).

### *hrcN^G176A^*-dysregulated effectors contribute to the onset of infection phenotypes

To determine the impact of reduced effector expression on *Arabidopsis* infection, *hopAA1-2, hopAM1, hopAF1,* and *hopH1* were expressed *in trans* in WT *Pto* and *hrcN^G176A^* and their impact on infection phenotypes was assessed (Fig 5). Overexpression of *hopAA1-2, hopAM1* and *hopAF1* had little or no impact on infection phenotypes (Fig 5A) or bacterial load (S4A-B Fig) in WT *Pto*, but led to a partial recovery of chlorosis and necrosis in *hrcN^G176A^*. Conversely, overexpression of *hopH1* had little effect on infection phenotypes with either tested strain.

**Fig 5A-B:**
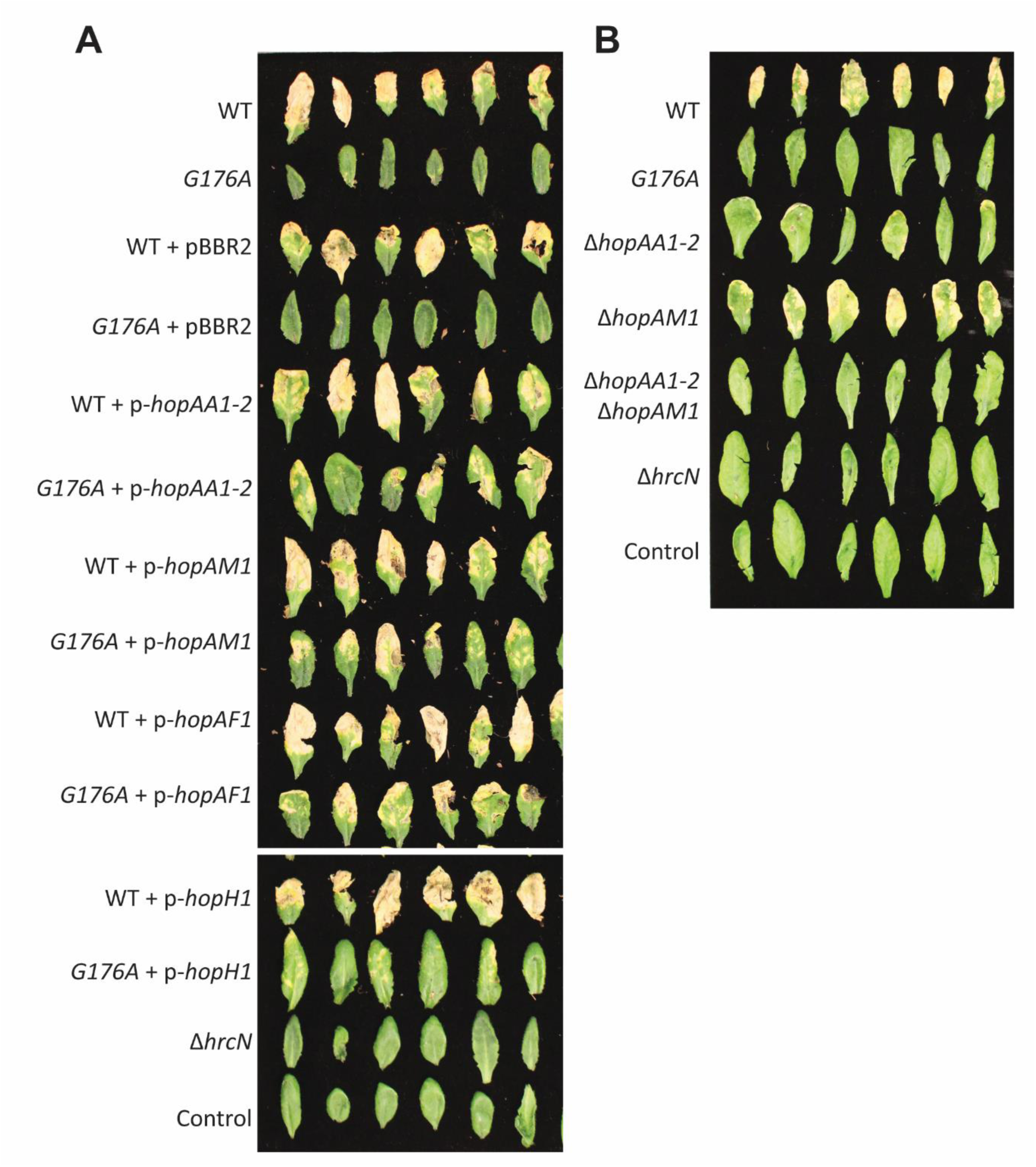
Visual disease phenotypes of *Pto* DC3000 infiltrated *A. thaliana* Col-0 leaves, 6 days post- infection. **A:** *Pto* strains harbour the *hrcN^G176A^* allele/Δ*hrcN* and effector expression plasmids as indicated. pBBR2: empty vector. **B:** *Pto* strains harbour the *hrcN^G176A^* allele or effector/*hrcN* deletions as indicated. In each case, control indicates uninfected leaves and leaves were taken at random from three independent plants. *ti.hopAM1*

Finally, we examined the impact of effector deletion mutants on Col0 infections. While our efforts to generate Δ*hopAF1* were unsuccessful, we were able to produce *Pto* Δ*hopAA1-2*, Δ*hopAM1* and a Δ*hopAA1-2/hopAM1* double mutant. Strikingly, while *hopAM1* deletion produced no obvious effects on *Pto* infection phenotypes, both *hopAA1-2* deletion mutants phenocopied the *hrcN^G176A^* strain, with a loss of infection phenotypes (Fig 5B) but no accompanying reduction in bacterial load (S4C-D Fig). Together, these data strongly suggest a central role for HopAA1-2 in enabling symptom onset during *Pto Arabidopsis* infections.

## Discussion

Effective plant infection by phytopathogenic bacteria relies on the coordinated deployment of T3Es (6, 38). In addition to regulating T3I deployment and activity (10, 45), and controlling the delivery of specific T3Es to the injectisome (44), our findings suggest that secretion of specific T3Es is also controlled by allosteric regulation of T3I function by the second messenger cdG. Conservative point mutations in residues linked to *hrcN* cdG binding (54) led to markedly altered virulence phenotypes upon *A. thaliana* infection, with *Pto hrcN^G176A^* and *hrcN^G311A^* mutants showing few visible signs of disease, despite bacterial apoplastic proliferation remaining unaffected.

The asymptomatic infection phenotypes seen for the *Pto hrcN^G176A/G311A^* mutants were not due to compromised T3I assembly, or a nonspecific reduction in effector secretion. The T3I apparently assembles normally in both mutants, and ATPase activity and cdG binding remain intact in HrcN^G176A^. However, while secretion of many T3Es is unaffected relative to wild-type *Pto*, the *hrcN^G176A/G311A^* mutants show disrupted export of a specific subset of T3Es, including several responsible for the onset of disease symptoms such as tissue chlorosis and necrosis.

Immunocompromised plants showed wild-type infection symptoms upon infiltration with both *hrcN* mutants, suggesting that the disrupted effectors function by suppressing elements of the plant immune response, which in-turn counter the destruction of plant tissues during infection. Furthermore, wild- type infection symptoms were recovered in both mutants by manipulating intracellular cdG levels, suggesting that they represent either a reduction in HrcN nucleotide binding affinity, or a partial decoupling of the ability to bind cdG from the downstream impact of dinucleotide binding on export apparatus function. Further research is needed to fully dissect the relationship between cdG and HrcN complex formation in *P. syringae*.

The decoupling we observe between leaf disease symptoms and bacterial proliferation in *P. syringae* infections appears to be a rare phenomenon, with changes to disease symptom severity typically coupled to a corresponding increase or decrease in apoplastic colonisation (59, 60). Symptomless *P. syringae* infections have previously been reported where the induction of HR in non-host plants was missing (61). Decoupling of disease severity and colonisation has also been reported for mutants in phytotoxin pathways. *Pto* strains lacking syringolin A undergo asymptomatic infections of wheat plants, whereas bacterial colonisation is relatively unaffected (62). A similar reduction in necrotic lesion formation has been also reported for strains missing the phytotoxin coronatine, albeit in this case accompanied by a slight drop in bacterial load (59).

HopAM1, HopAF1 and particularly HopAA1-2 appear to play key roles in mediating *Pto* symptom development, and their export is post-transcriptionally regulated by HrcN. Using ELISA analysis of T3E- CyaA reporter fusion strains, we showed that delivery of specific effector proteins into leaves, including HopAA1-2, HopAM1, and HopAF1 was significantly compromised in the *hrcN^G176A/G311A^* mutants relative to wild-type. While over-expression of all three effectors partially recovered disease phenotypes in infections with the asymptomatic *Pto hrcN^G176^* mutant, only *hopAA1-2* appeared to be necessary for full establishment of disease phenotypes, with *hopAM1* deletion mutants showing similar visual disease symptoms to wild-type *Pto*.

HopAA1-2, HopAM1 and HopAF1 have all previously been linked to immune system dysregulation, which aligns with our observation of symptom recovery for *hrcN^G176A^* infections in immune compromised plants. HopAA1-2 interacts with EDS1 (enhanced disease susceptibility 1) and PBS3, and is hypothesised to play a role in salicylic acid-mediated defence subversion (63). EDS1 is involved in transcriptional mobilisation associated with resistance pathways (64), while PBS3 protects EDS1 from proteasome-mediated degradation (65). Reduced levels of both genes were seen upon co-expression with HopAA1-2 (63). Deletion of a gene cluster containing *hopAA1-2* (alongside *hopV1*, *hopAO1, and hopG1*) showed strongly reduced *Pto* virulence in *N. benthamiana* and *A. thaliana* plants (66).

HopAM1 shares high Toll/interleukin-1 receptor (TIR) domain homology to NB-LRR receptors, which can hydrolyse nicotinamide adenine dinucleotide (NAD^+^) (67). It has been shown to induce meristem chlorosis and local cell death in *Arabidopsis* (68). EDS1, alongside HSP90.2 and SGT1b; part of a steady- state NLR chaperone complex were shown by genome-wide association mapping to affect HopAM1- induced cell death (68). HopAM1 has also been linked to virulence enhancement in water-stressed plants by inducing hypersensitivity to abscisic acid (69).

Finally, HopAF1 can suppress plant immunity by blocking ethylene induction through targeting of methionine recycling (70). HopAF1 showed an interaction with Arabidopsis methylthioadenosine nucleosidase (MTN1 and MTN2) in yeast 2-hybrid screens (70, 71). MTN enzymes are known to be involved in the Yang cycle, a process essential for high levels of cellular ethylene in *A. thaliana* (70). HopAF1 has also been shown to suppress production of defensive reactive oxygen species in the phytopathogen *Pseudomonas savastanoi* (72).

Bacterial cdG signalling relays diverse environmental inputs into integrated phenotypic responses (48). This control can take place both at the level of transcription, but also by rapidly influencing the activity of existing proteins in the cell, as seen here. We propose that cdG binding to HrcN enables the allosteric control of T3E export for a subset of effectors involved in disease symptom development. Our model suggests that the coordinated deployment of bacterial virulence factors depends not only on transcriptional hierarchy and bacterial proximity to apoplastic cells, but is controlled actively and rapidly in response to signals from the apoplastic environment.

## Materials and Methods

### Bacterial strains and growth conditions

Unless otherwise stated *P. syringae* strains were grown at 28°C and *E. coli* at 37°C with shaking. Bacterial strains and plasmids used in this study are listed in Table S1. King’s B medium (KB) (73) and L medium (74) were used for bacterial cell culture as specified. Antibiotics were used at final concentrations of rifampicin at 50 μg/ml, nystatin at 25 µg/mL, tetracycline at 12.5 μg/ml, kanamycin at 50 μg/mL and chloramphenicol at 25 μg/mL, unless otherwise stated.

### Molecular cloning

Cloning was performed following standard molecular biology techniques (74). *Pto* gene deletions and *hrcN* mutants were generated by allelic exchange, using vectors constructed by amplifying the upstream and downstream flanking regions of each mutagenesis site with primers DC3000-HrcN-FWD and DC3000-HrcN-REV plus appropriate internal point mutant primers (e.g., *Mut*-FWD/REV-DC3000- HrcN, see Table S2), combining them by ligation/strand overlap extension PCR as appropriate and cloning into the suicide vector pTS1 between restriction sites *Xho*I and *Bam*HI (75). *Pto* cells were transformed by electroporation with the appropriate deletion/mutagenesis vectors following the method described in (76). Single crossover integrations of each plasmid into the chromosome were selected on tetracycline plates and re-streaked before single colonies were grown overnight in KB media without selection. Following a tetracycline enrichment step after (77), double crossovers were counter-selected by plating serial dilutions onto L agar containing 10% (w/v) sucrose. Deletion/site- specific *hrcN* mutants were subsequently confirmed by PCR and sequencing using primers FWDExternalHrcN and REVExternalHrcN (Table S2).

To complement *Pto hrcN/hrcC* mutations with WT gene copies, the *hrcN/hrcC* genes and ∼200 bp of upstream DNA incorporating the promoter region were amplified using the HrcNComp FWD and REV primers (Table S2) and cloned into pUC18-miniTn7-Gm between *Hind*III and *Bam*HI/*Pst*I. The resulting vectors were transformed into *Pto* mutant strains by electroporation alongside the helper plasmid pTNS2 (78). Chromosomal integration at the *att*::Tn7 site for successful transformants was verified by PCR using primers pTN7R and pGlmS-Down (78). HrcN purification vectors were constructed by amplifying the *hrcN* alleles synthesised above using primers OE_DC3000_FWD and OE_DC3000_REV plus relevant internal point mutant primers (Table S2) and cloning the resulting fragments between the *NdeI* and *XhoI* restriction sites of a pETM11 expression vector. The effector-CyaA fusion constructs were produced by gateway cloning of effectors and plasmids, as described previously (58). pCPP5371- CyaA backbone vectors were Gateway cloned with T3E (lacking stop-codons)-pENTR-SD/D-TOPO vectors via an LR reaction to create pCPP5371::T3E + C-terminal CyaA vectors (pDEST) (58). Effector protein overexpression vectors were produced by amplifying effector genes using primers e.g. OvExp HopAF1 FWD/REV (see Table S2 for complete list) before cloning the resulting fragments between the *Kpn*I *and Xho*I restriction sites of a pBBR1MCS-2 expression plasmid (79).

### Protein purification

Protein purification was conducted after (77). Briefly, 5 mL overnight cultures of *E. coli* BL21-(DE3) pLysS containing pETM11-*hrcN*-His_6_ vectors were used to inoculate 1.0 litre terrific broth overexpression cultures. These were then grown at 37°C, 220 rpm to an OD_600_ of 0.4, before protein expression was induced overnight at 18°C, 220 rpm with 1mM IPTG. Cells were harvested at 5000 g, 4°C for 10 minutes and re-suspended in 30 mL of ice cold chromatography binding buffer (20 mM HEPES, 250 mM NaCl, 2 mM MgCl2, 3 % glycerol pH 7.8), before immediate lysis on ice using an MSE Soniprep 150 ultrasonic disintegrator. The soluble fraction was loaded onto a HisTrap excel (Cytiva) column equilibrated with binding buffer. Following protein immobilisation, a column wash with 10% elution buffer (20 mM HEPES, 250 mM NaCl, 2 mM MgCl_2_, 3% glycerol, 500 mM imidazole pH 7.8) was performed to remove non-specific contaminants. Proteins were eluted over a linear gradient of 10- 100% elution buffer. Protein purity was confirmed by SDS-PAGE and concentrations were calculated using Bradford assays (80). Purified HrcN proteins were taken forward immediately for *in vitro* analysis.

### Protein structural predictions

The hexameric structure of HrcN was modelled by AlphaFold3 (81), using the default settings and specifying 6 monomeric subunits. The downstream analysis and image generation was performed in Pymol.

### Linked pyruvate kinase / lactate dehydrogenase (PK/LDH) ATPase activity assays

ATPase activity was measured indirectly by monitoring NADH oxidation in a FLUOstar Omega or a SPECTROstar Nano plate reader (BMG Labtech) at 25 °C. The reaction buffer consisted of 100 mM Tris- Cl (pH 9.0), 20 mM MgSO4. Each 100 μL reaction contained 1 μM (final concentration) of purified HrcN, 0.4 mM NADH, 0.8 mM phosphoenolpyruvic acid, 0.7 μL PK/LDH enzyme (Sigma) and was initiated by the addition of 10 μL ATP solutions, 2 replicates of each concentration: 0, 0.125, 0.25, 0.5, 0.75, 1.0, 1.5, 2.0 mM ATP. Enzyme kinetics were determined by measuring A_340_ at 1-minute intervals. Kinetic parameters were calculated by plotting the specific activity of the enzyme (nmol ATP hydrolyzed/min/mg protein) versus [ATP] and by fitting the non-linear enzyme kinetics model (Michaelis-Menten) in GraphPad Prism.

### DRaCALA nucleotide binding assays

CdG/CdA binding assays were performed as described by Roelofs et al. (82). Protein samples were incubated for 2 minutes at room temperature with 2’-Fluo-AHC-c-diGMP (BioLog 009) or 2’-Fluo-AHC- c-diAMP (BioLog C 088), in a final concentration of 0.6 μΜ in reaction buffer (25 mM Tris, 250 mM NaCl, 10 mM MgCl2, 5 mM β-mercaptoethanol pH 7.5). For the competition experiments, 1 mM of each competing nucleotide was mixed with 1.25 μΜ HrcN and incubated for 30 minutes prior to the addition of the fluorescent di-nucleotides. The tubes were incubated on ice for 10 minutes in the dark. 5 μL of each sample was then spotted on nitrocellulose membrane and left to briefly dry before visualisation using the UV fluorescence mode on a G:Box F3 Imager (Syngene).

### Plant growth conditions

Col0 *A. thaliana* plants and immuno-compromised fec (*fls2, efr,* and *cerk1* gene knock-out) and bbc (*bak1-5, bkk1-1,* and *cerk1* gene knock-out) lines (83) were used in this study. Plants were grown in controlled environment horticultural facilities under short day (10-hour) lighting conditions [120-180 μmol m^-2^ s^-2^ light intensity], 70 percent relative humidity, and 22°C growth temperature. All seedlings were grown prior to infection for 4-5 weeks + 1 week 4°C seed vernalisation in mixed *Arabidopsis* peat (Levington F2 600 ITS Peat, 100 ITS 4mm grit, 196g Exemptor Chloronicotinyl Insecticide). Plant experiments were conducted in containment level 2 controlled environment rooms, under plant health licence 51103/197332/5.

### Plant infection assays

Cell cultures were grown overnight in L medium, harvested by centrifugation, then resuspended in 10mM MgCl_2_ and adjusted to a final density (OD_600_) of 0.0002 (equivalent to 10^7^ cells ml^-1^). Cells were gently infiltrated into the leaf apoplast using a 1 ml syringe until leaves appeared wet. Post infiltration, bacterial suspensions were allowed to absorb/dry onto leaf surfaces for approximately 1 h before initial (0D) samples were collected. For calculating bacterial load, two 7mm diameter leaf discs (area 0.384 cm^2^) were collected for each sample at 0, 2 and 3-days post-infection, flash-frozen, and ground using a Qiagen Tissue Lyser II (30 hz, 45 seconds x 2). Lysate was serially diluted 10^-1^ to 10^-5^ and plated onto LB with rifampicin and nystatin. Plates were incubated for 2 days at 28 °C prior to colony counting. Bacterial load was calculated and presented as colony forming units per unit leaf area (CFU/cm^2^). A minimum of eight plants were used for each condition and assays were run at least twice independently.

### Western blotting of leaf tissue

Two 7mm discs of *Pto*-infected leaf tissue per sampled plant (3 days post-infection) were incubated with 100 µl SDS-PAGE sample buffer (BioRad) at 100°C for 10 minutes, before the supernatant was collected following centrifugation (16,000 g, 10 minutes, 4°C) to pellet the insoluble plant material. Samples were then separated by SDS-PAGE gels and transferred onto a polyvinylidene difluoride (PVDF) membrane (Millipore). Membranes were incubated overnight in blocking solution at 8°C (1x PBS pH 7.4, 0.01% Tween-20, 5% milk powder). Protein was subsequently detected with 1/10,000 anti-HrcN/HrcC primary antisera (Minotech Biotechnology) and 1/3000 anti-rabbit secondary antibody (Sigma). Western blots were visualized using ECL chemiluminescent detection reagent on an ImageQuant LAS- 500 scanner (GE Healthcare).

### ImageJ analysis

The level of chlorosis severity of infected leaves was quantified by average pixel analysis using ImageJ (84). Images were opened and the leaf surface area was selected using the multi-point tool for each leaf. The measure setting was used to calculate pixel intensity (brightness of a given pixel) over the selected area on a scale of 1 to 255 (1 showing least pixel intensity and no evidence of chlorosis, 255 showing highest pixel intensity with strong evidence of chlorosis). A minimum of 8 leaves per sample were used to calculate final averages per replicate.

### CyaA effector fusion reporter assay

Pto strains containing effector-CyaA fusion constructs (in pCPP5295 stable expression vectors, driven by a *hrp*-promoter) were grown overnight in KB medium and diluted to an OD_600_ of 0.05 (5 × 10^7^ cfu/mL) in 10 mM MgCl_2_ before infiltration into 4-week-old Col0 leaves as described above. 6 hours post- infection, 4 mm diameter leaf discs from the infected leaves were collected using a leaf corer, flash frozen in liquid nitrogen and ground using a Qiagen Tissue Lyser II (30 hz, 45 seconds x 2). 300 µL of 0.1 M HCl was added to each sample, samples were pelleted by centrifugation (13,000 g, 10 min), vortexed and then pelleted by centrifugation for a further 10 minutes. The supernatant was collected and diluted 1:5 and 1:50 in 0.1 M HCl. A direct cAMP ELISA kit (Enzo) was used to quantify [cAMP] in each sample following the manufacturer’s instructions. Data was analysed by comparing each well against the logarithmic curve generated from the cAMP standards. This value was then divided by the calculated sample protein concentration (via standard Bradford assay) to give pmol cAMP/µg protein.

## Acknowledgements

The authors would like to thank Eleftheria Trampari and Isabella Frost for insightful conversations about the project, and Lucia Grenga and Sebastian Pfeilmeier respectively for the pBBR4-*wspR19* and pBBR4- *bifA* plasmids. JGM, CMAT and RHL were funded by BBSRC Institute Strategic Programme grants BBS/E/J/000PR9797 and BB/X010996/1 to the John Innes Centre. DW was funded by a BBSRC Doctoral Training Partnership (BB/J014524/1) PhD studentship.

---

**S1 Table:** Strains and Plasmids used in this study.

| Strain Name | Genotype | Reference |
| --- | --- | --- |
| ***Escherichia coli*** |  |  |
| DH5α | endA1, hsdR17(rK-mK+), supE44, recA1, gyrA (Nal<sup>r</sup>), relA1, Δ(lacIZYA-argF) U169, deoR, Φ80dlacΔ(lacZ)M15 | (85) |
| DB3.1 | gyrA462 endA1 Δ(sr1-recA) mcrB mrr hsdS20 glnV44 (=supE44) ara14 galK2 lacY1 proA2 rpsL20 xyl5 leuB6 mtl1 | Invitrogen |
| BL21-(DE3) pLysS | Sm<sup>R</sup>, K12 recF143 lacIq lacZΔ.M15, xylA, pLysS | Novagen |
| ***Pseudomonas syringae* pv *tomato* DC3000 (Pto)** |  |  |
| WT Pto DC3000 | Rif<sup>R</sup> derivative of *P. syringae* pv *tomato* NCPPB 1106 | (86) |
| *Pto hrcN*<sup>G176A</sup> | WT Pto DC3000 with a G176A mutation in *hrcN* | This study |
| *Pto hrcN*<sup>E208D</sup> | WT Pto DC3000 with an E208D mutation in *hrcN* | This study |
| *Pto hrcN*<sup>G311A</sup> | WT Pto DC3000 with a G311A mutation in *hrcN* | This study |
| *Pto hrcN*<sup>L338V</sup> | WT Pto DC3000 with an L338V mutation in *hrcN* | This study |
| *Pto hrcN*<sup>F174Y</sup> | WT Pto DC3000 with an F174Y mutation in *hrcN* | This study |
| *Pto ΔhrcC* | WT Pto DC3000 with an inactivating mutation in *hrcC* | (87) |
| *Pto ΔhrcN* | WT Pto DC3000 with a non-polar deletion of *hrcN* | This study |
| *Pto ΔhopAA1-2* | WT Pto DC3000 with a non-polar deletion of *hopAA1-2* | This study |
| *Pto ΔhopAM1* | WT Pto DC3000 with a non-polar deletion of *hopAM1* | This study |
| *Pto ΔhopAF1* | WT Pto DC3000 with a non-polar deletion of *hopAF1* | This study |
| *Pto ΔhopAA1-2 ΔhopAM1* | WT Pto DC3000 with non-polar deletions of *hopAA1-2* & *hopAM1* | This study |
| *Pto* WT + *hrcC* | WT Pto DC3000 with *hrcC* inserted at the *att::Tn7* site. Gent<sup>R</sup> | This study |
| *Pto* WT + *hrcN* | WT Pto DC3000 with WT *hrcN* inserted at the *att::Tn7* site. Gent<sup>R</sup> | This study |
| *Pto ΔhrcC* + *hrcC* | *Pto ΔhrcC* with *hrcC* inserted at the *att::Tn7* site. Gent<sup>R</sup> | This study |
| *Pto ΔhrcN* + *hrcN* | *Pto ΔhrcN* with WT *hrcN* inserted at the *att::Tn7* site. Gent<sup>R</sup> | This study |
| *Pto hrcN*<sup>G176A</sup> + *hrcN* | *Pto hrcN*<sup>G176A</sup> with WT *hrcN* inserted at the *att::Tn7* site. Gent<sup>R</sup> | This study |
| *Pto hrcN*<sup>E208D</sup> + *hrcN* | *Pto hrcN*<sup>E208D</sup> with WT *hrcN* inserted at the *att::Tn7* site. Gent<sup>R</sup> | This study |
| *Pto hrcN*<sup>G311A</sup> + *hrcN* | *Pto hrcN*<sup>G311A</sup> with WT *hrcN* inserted at the *att::Tn7* site. Gent<sup>R</sup> | This study |
| **Plasmids** |  |  |
| pUC18-mini-Tn7-Gm | Amp<sup>R</sup>, Gent<sup>R</sup>, *att::Tn7* insertion vector | (88) |
| pUC18-mini-Tn7-*hrcN/hrcC* | pUC18-miniTn7-GM containing WT *hrcN/hrcC* cloned between *HindIII* and *BamHI/PstI* sites | This study |
| pTNS2 | Helper plasmid for *att::Tn7* insertion. Amp<sup>R</sup> | (88) |
| pTS1 | Tet<sup>R</sup>, suicide vector; *ColE1*-replicon, *IncP-1*, *Mob*, *lacZ* | (75) |
| pTS1- mutagenesis vectors | pTS1 containing various *hrcN* alleles/gene deletion constructs cloned between *XhoI* and *BamHI* sites | This study |
| pBBR1MCS-4 | Broad host-range expression vector. MCS with *lacZ* blue/white selection and upstream *lac* promoter. Amp<sup>R</sup> | (79) |
| pBBR4-*bifA* | pBBR1MCS-4 expressing *Pseudomonas fluorescens* SBW25 *bifA* | S. Pfeilmeier, unpublished |
| pBBR4-*wspr19* | pBBR1MCS-4 expressing the *P. fluorescens* SBW25 *wspr19* allele | L. Grenga, unpublished |
| pETM11 | Protein expression vector, enabling N-terminal His<sub>6</sub> tag fusions. Kan<sup>R</sup> | (89) |
| pETM11-*hrcN* vectors | pETM11 with *hrcN* alleles cloned between *NdeI* and *XhoI* sites | This study |
| pBBR1MCS-2 | Broad host-range expression vector. MCS with *lacZ* blue/white selection and upstream *lac* promoter. Kan<sup>R</sup> | (79) |
| pBBR2-*hopAA1-2* | pBBR1MCS-2 with *hopAA1-2* cloned between *KpnI* and *XhoI* sites | This study |
| pBBR2-*hopAM1* | pBBR1MCS-2 with *hopAM1* cloned between *KpnI* and *XhoI* sites | This study |
| pBBR2-*hopAF1* | pBBR1MCS-2 with *hopAF1* cloned between *KpnI* and *XhoI* sites | This study |
| pBBR2-*hopH1* | pBBR1MCS-2 with *hopH1* cloned between *KpnI* and *XhoI* sites | This study |
| pCPP5371 | Gateway destination vector for expression of *Pto* effectors from the *avrPto* promoter with C-terminal *cya* tags. Gent<sup>R</sup>, Cm<sup>R</sup> | (90) |
| pENTR-SD/D-TOPO vectors | pENTR-SD/D-TOPO gateway cloning vectors containing *Pto* effector genes lacking stop codons. Kan<sup>R</sup> | (91) |
| pDEST vectors | pCPP5371 vectors containing *Pto* effector genes with C-terminal *cya* tags. Gent<sup>R</sup>, Cm<sup>R</sup> | This study |

**S2 Table:**
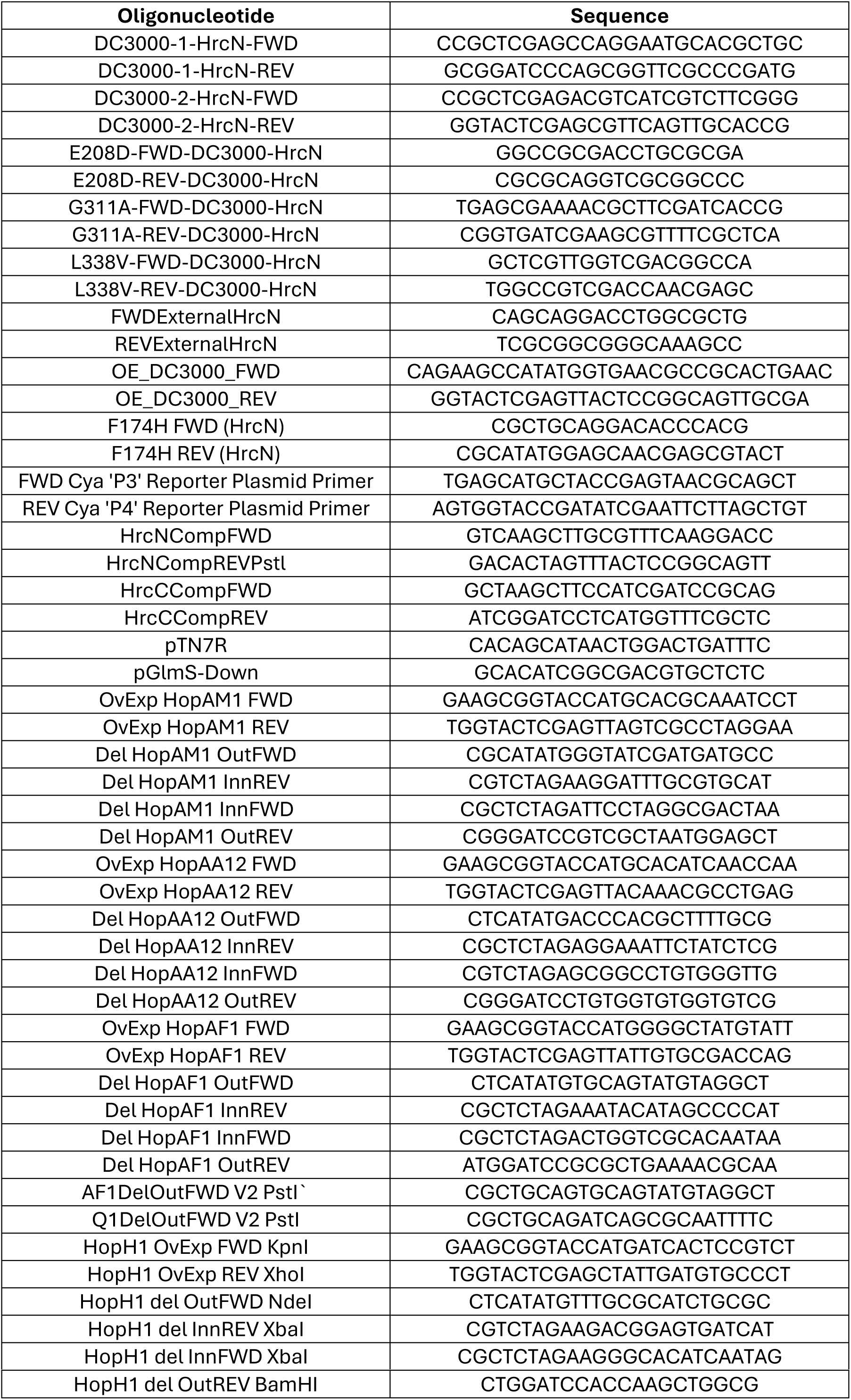
Primers used in this study.

## Supplementary Figure Legends

**S1A Fig:**
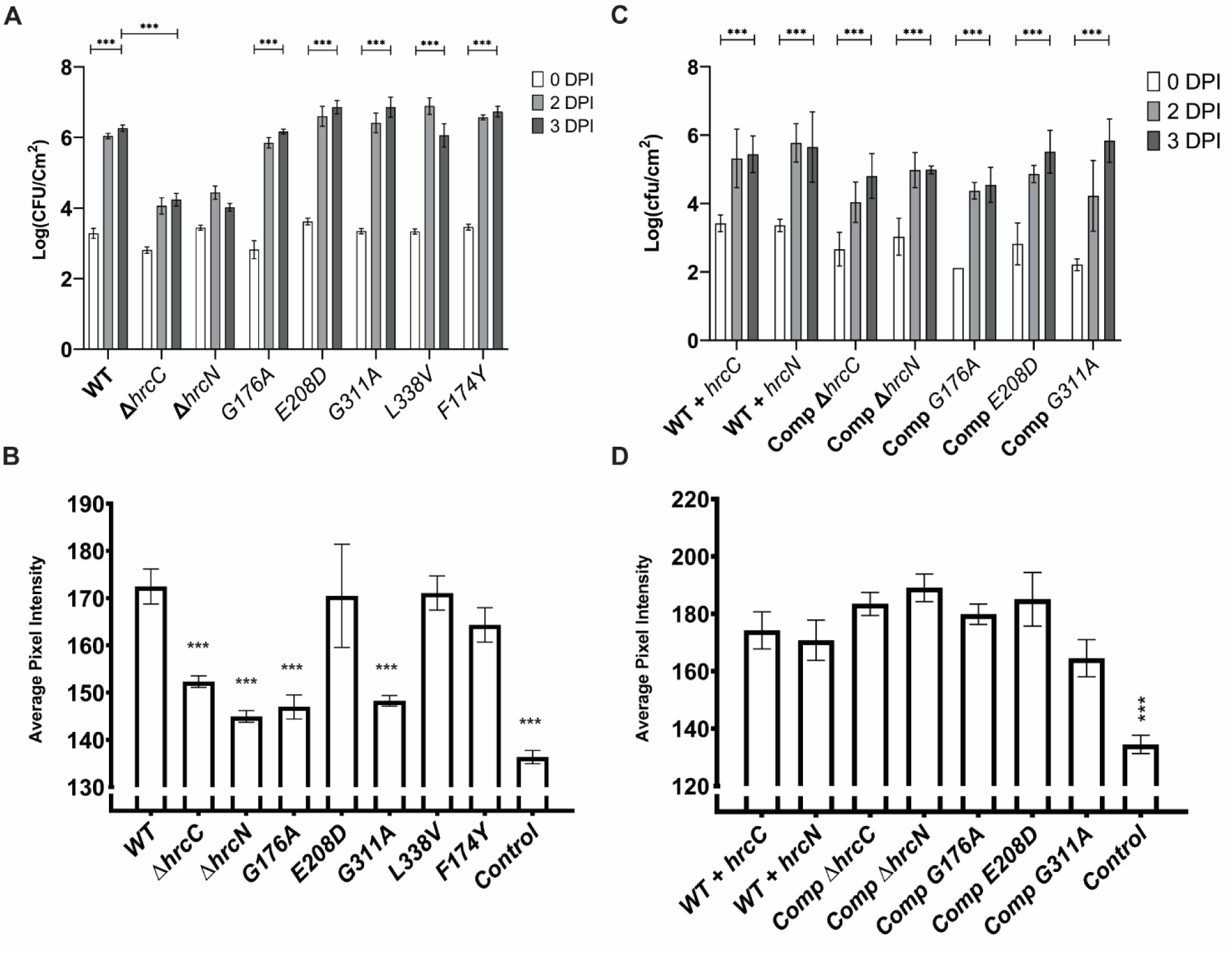
Colony forming units (CFU) recovered from *A. thaliana* Col-0 leaves infiltrated with *Pto* DC3000 *hrcC/hrcN* mutants at 0, 2 and 3 DPI as indicated. **S1B:** Average pixel intensity analysis for *A. thaliana* Col-0 leaves infiltrated with *Pto* DC3000 *hrcC/hrcN* mutants 6 days post-infection. Analysis was performed using ImageJ software (version 1.52a) and increased intensity is directly proportional to the extent of leaf yellowing. Control indicates uninfected leaf tissue. **S1C:** Colony forming units (CFU) recovered from *A. thaliana* Col-0 leaves infiltrated with *Pto* DC3000 *hrcC/hrcN* complementation strains at 0, 2 and 3 DPI as indicated. **S1D:** Average pixel intensity analysis for *A. thaliana* Col-0 leaves infiltrated with *Pto* DC3000 complementation strains 6 days post-infection. Analysis was performed using ImageJ software (version 1.52a) and increased intensity is directly proportional to the extent of leaf yellowing. Control indicates uninfected leaf tissue. In each case, different *hrcC/hrcN* alleles are indicated on the X-axis. Error bars show standard error of the mean, and asterisks denote statistically significant differences from the WT/D0 (2 sample t-test) where ‘***’ denotes p = ≤ 0.001. **A:** n = 4 plants; **C:** n = 3 plants; **B,D:** n = 8 leaves.

**S2A Fig:**
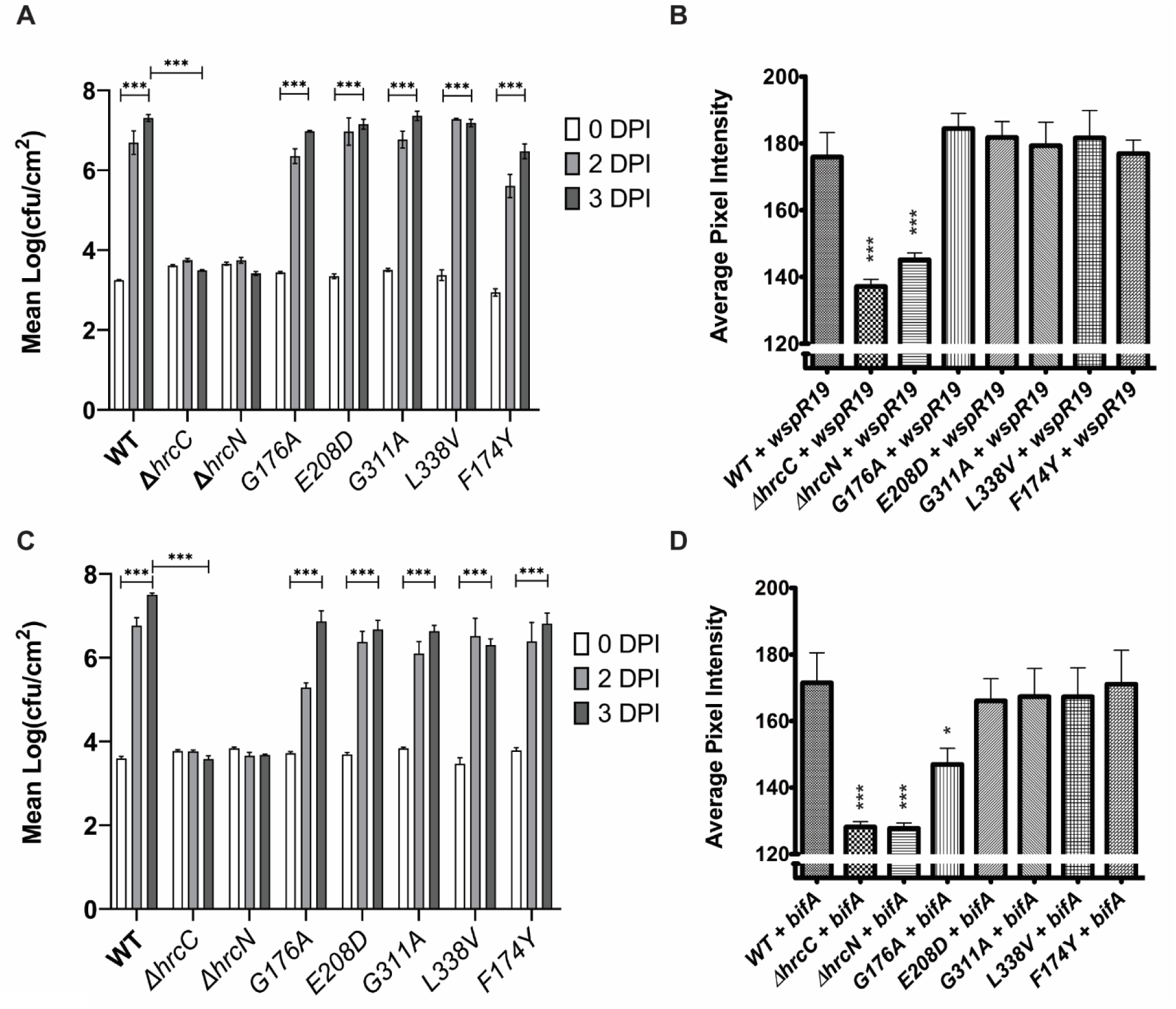
Colony forming units (CFU) recovered from *A. thaliana* Col-0 leaves infiltrated with *Pto* DC3000 *wspR19-*expressing strains at 0, 2 and 3 DPI as indicated. **S2B:** Average pixel intensity analysis for *A. thaliana* Col-0 leaves infiltrated with *Pto* DC3000 *wspR19-*expressing strains 6 days post-infection. Analysis was performed using ImageJ software (version 1.52a) and increased intensity is directly proportional to the extent of leaf yellowing. **S2C:** Colony forming units (CFU) recovered from *A. thaliana* Col-0 leaves infiltrated with *Pto* DC3000 *bifA-*expressing strains at 0, 2 and 3 DPI as indicated. **S2D:** Average pixel intensity analysis for *A. thaliana* Col-0 leaves infiltrated with *Pto* DC3000 *bifA-*expressing strains 6 days post-infection. Analysis was performed using ImageJ software (version 1.52a) and increased intensity is directly proportional to the extent of leaf yellowing. In each case, different *hrcC/hrcN* alleles are indicated on the X-axis. Error bars show standard error of the mean, and asterisks denote statistically significant differences from the WT/D0 (2 sample t-test) where ‘*’ = p ≤ 0.05 and ‘***’ = p ≤ 0.001. **A,C:** n = 3 plants; **B,D:** n = 8 leaves.

**S3A Fig:**
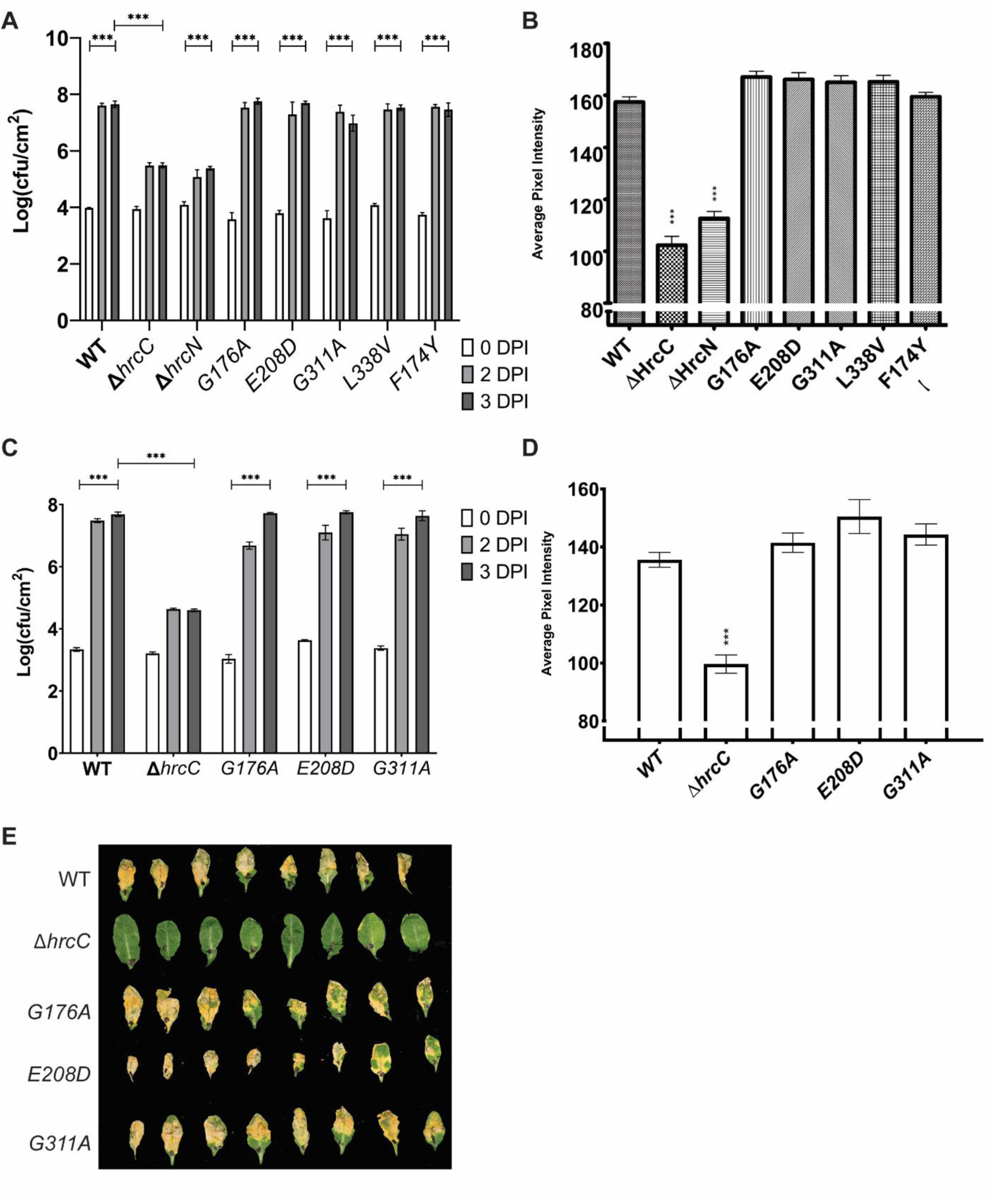
Colony forming units (CFU) recovered from *A. thaliana bbc* (immunocompromised) leaves infiltrated with *Pto* DC3000 *hrcC/hrcN* mutants at 0, 2 and 3 DPI as indicated. **S3B:** Average pixel intensity analysis for *A. thaliana bbc* leaves infiltrated with *Pto* DC3000 *hrcC/hrcN* mutants 6 days post- infection. Analysis was performed using ImageJ software (version 1.52a) and increased intensity is directly proportional to the extent of leaf yellowing. **S3C:** Colony forming units (CFU) recovered from *A. thaliana fec* leaves infiltrated with *Pto* DC3000 *hrcC/hrcN* mutants at 0, 2 and 3 DPI as indicated. **S3D:** Average pixel intensity analysis for *A. thaliana fec* (immunocompromised) leaves infiltrated with *Pto* DC3000 *hrcC/hrcN* mutants 6 days post-infection. Analysis was performed using ImageJ software (version 1.52a) and increased intensity is directly proportional to the extent of leaf yellowing. In each case, different *hrcC/hrcN* alleles are indicated on the X-axis. Error bars show standard error of the mean, and asterisks denote statistically significant differences from the WT (2 sample t-test) where ‘***’ denotes p = ≤ 0.001. **A,C:** n = 3 plants; **B,D:** n = 8 leaves. **S3E:** Visual disease phenotypes of *Pto* DC3000 infiltrated *A. thaliana fec* leaves taken at random from 3 independent plants, 6 days post- infection. *Pto* strains harbour the *hrcC/hrcN* alleles as indicated.

**S4A Fig:**
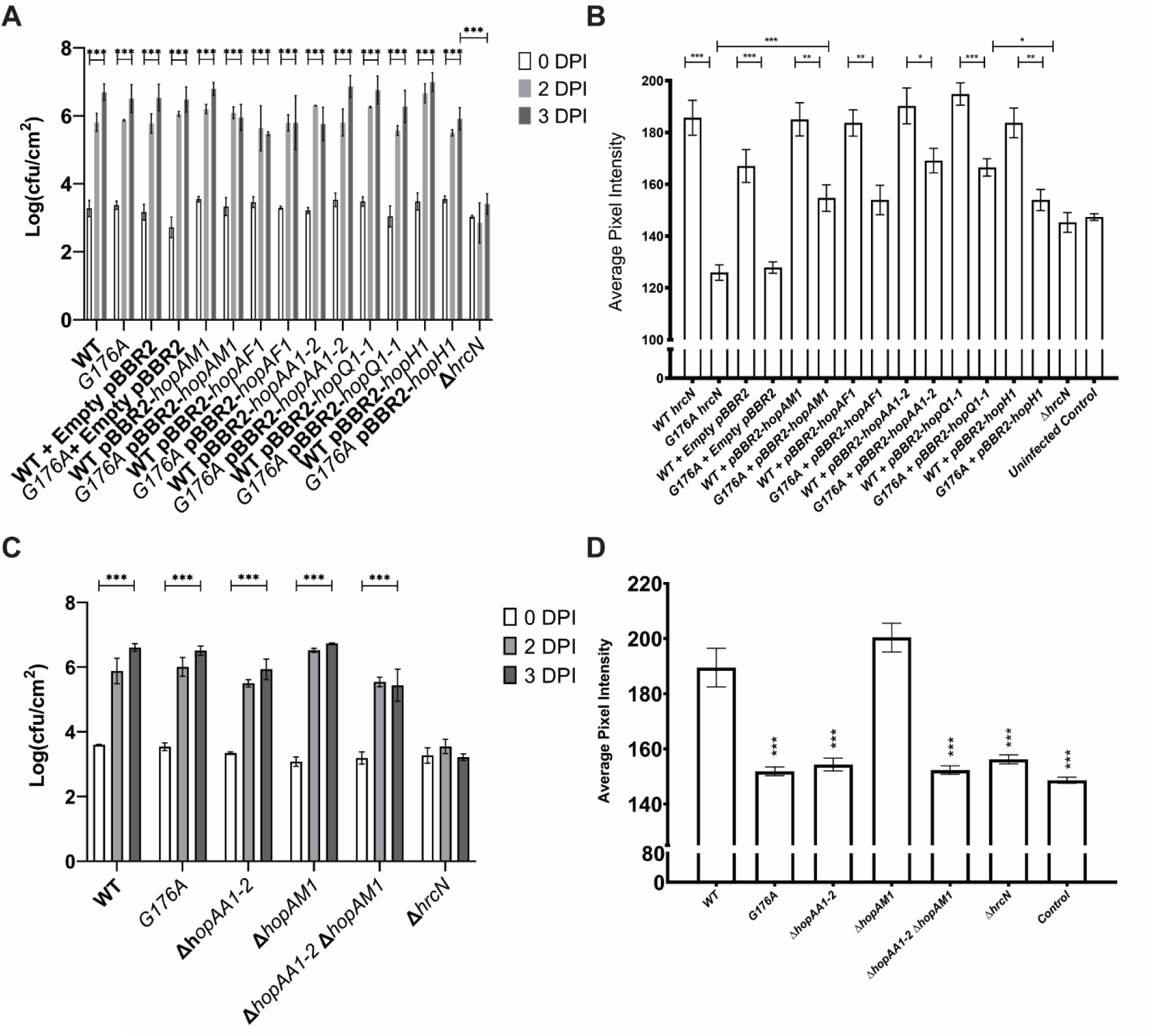
Colony forming units (CFU) recovered from *A. thaliana* Col-0 leaves infiltrated with *Pto* DC3000 effector overexpression strains at 0, 2 and 3 DPI as indicated. **S4B:** Average pixel intensity analysis for *A. thaliana* Col-0 leaves infiltrated with *Pto* DC3000 effector overexpression strains 6 days post- infection. Analysis was performed using ImageJ software (version 1.52a) and increased intensity is directly proportional to the extent of leaf yellowing. Control indicates uninfected leaf tissue. **S4C:** Colony forming units (CFU) recovered from *A. thaliana* Col-0 leaves infiltrated with *Pto* DC3000 effector deletion mutants at 0, 2 and 3 DPI as indicated. **S4D:** Average pixel intensity analysis for *A. thaliana* Col- 0 leaves infiltrated with *Pto* DC3000 effector deletion mutants 6 days post-infection. Analysis was performed using ImageJ software (version 1.52a) and increased intensity is directly proportional to the extent of leaf yellowing. Control indicates uninfected leaf tissue. In each case, different *hrcN/*effector mutants are indicated on the X-axis. Error bars show standard error of the mean, and asterisks denote statistically significant differences from the WT/D0 (2 sample t-test) where ‘*’ = p ≤ 0.05, ‘**’ = p ≤ 0.01, and ‘***’ = p ≤ 0.001. **A,C:** n = 3 plants; **B,D:** n = 8 leaves.

